# In Vitro Fertilization using Magnetotactic Sperm Cells

**DOI:** 10.64898/2026.04.23.720095

**Authors:** Carla Ribeiro, Friedrich Striggow, Richard Nauber, Franziska Hebenstreit, Jennifer Schoen, Mariana Medina-Sánchez

## Abstract

In vitro fertilization (IVF) is essential for many couples facing infertility, e.g. in cases of low sperm count (oligospermia), where natural fertilization is unlikely. Medical microrobotics, making use of microscopic devices designed to perform targeted tasks inside the body under imaging guidance and controlled actuation, represents a promising strategy to guide sperm cells toward the oocyte. This approach may significantly reduce the time, invasiveness, and patient burden of conventional IVF, with long-term potential for in vivo assisted reproduction. Here, we report the first successful in vitro fertilization (IVF) using magnetically functionalized spermatozoa, termed magnetotactic sperm cells (MSCs), as a step toward in vivo microrobotic guidance of sperm cells for targeted artificial insemination. We present a protocol for the preparation of MSCs for their use in IVF, resulting in samples largely free of non-functionalized sperm cells (99.69% purity). We systematically evaluate the effect of particle functionalization on sperm health, including acrosome integrity, DNA fragmentation, mitochondrial membrane potential, oxidative stress, and epithelial interactions, and observe no adverse effects. Notably, MSCs showed improved mitochondrial membrane integrity compared to the control samples after two hours of incubation. Using MSCs, we successfully performed complete IVF cycles that resulted in embryos developing to the blastocyst stage at a comparable rate as non-functionalized sperm cells of the same concentration. Lower concentrations of non-functionalized sperm cells (comparable to those remaining in the MSC sample after purification) did not result in any development of embryos to blastocysts. To facilitate manipulation and translation, we implemented automated image-based recognition, magnetic manipulation, and pre-clustering routines that increased guidance efficiency and are compatible with standard IVF workflows. Together, these results demonstrate that magnetic functionalization can be applied without compromising key sperm quality metrics and can enable directed sperm guidance for assisted oocyte fertilization. This work provides a practical framework for integrating microrobotic sperm manipulation into assisted-reproduction workflows and supports further development toward automated in vitro and eventual in vivo applications.

## Introduction

Medical microrobots (MMRs) are miniature devices that can be actuated by external or local stimuli and are designed for diverse biomedical applications. They are promising tools for insitu diagnostics, local application of therapeutics, and tissue engineering^1,2^. MMRs hold promise in gynecology and reproductive medicine, including transport of gametes or zygotes within the female reproductive tract and targeted drug delivery for cancer therapy. ^1,3^ The focus of assisted reproductive techniques is to overcome infertility, the inability to achieve a pregnancy after 12 months of regular intercourse without the use of any contraceptive treatment.^4^ Current treatments for infertility include artificial insemination, in vitro fertilization (IVF), and intracytoplasmic sperm injection (ICSI), chosen according to the cause and severity of infertility.^5,6^ For male-factor infertility, characterized by reduced sperm motility (asthenozoospermia) or low sperm concentration (oligospermia),^7^ ICSI is commonly used.^8^ Even though IVF and ICSI are commonly and successfully employed in infertility treatment, embryo implantation rates remain low per embryo transfer (20-40 %).^8^ A potential alternative, particularly in cases of male factor infertility, involves the use of untethered microcarriers, comparable in size to sperm cells, designed to assist sperm in reaching the oocyte more efficiently. These microcarriers offer several potential advantages for both in vitro and in vivo assisted fertilization, including delivery of enzymes to overcome biological barriers, scavenging reactive oxygen species, protecting sperm during transport, improve imaging contrast while operating under tissue, carrying therapeutic cargos, and enabling in-situ sperm selection as previously reported by our group.^9–13^ However, these demonstrations have been limited to a small number of sperm cells or to answer fundamental research questions, including the study of sperm-microrobots in viscoelastic fluids observing flagellar dynamics, along with their drug loading capacity toward targeted delivery of drugs for diseases in the reproductive tract, including cancer.^13–16^ To increase the chances of fertilization or to control drug dosage in targeted drug delivery applications, it is necessary to transport multiple sperm cells.^10^ Various strategies for transporting multiple sperm cells have been considered, including multi-cell carrier structures^9^ or binding of sperm cells to hyaluronic acid microflakes,^10^ which are limited to a few tens to a few hundreds of sperm cells. Besides a large number of transported sperm cells, fast and simple coupling techniques and methods for sperm cell release are required, with robust magnetic control of multiple microrobots, either sequentially or as swarms.

By incorporating magnetic micro- or nanoparticles into sperm-hybrid microrobots, the complexity of the magnetic microstructures and of sperm-microstructure binding can be greatly reduced. This strategy has been used for the functionalization of immotile ^23^ as well as motile sperm cells, allowing their directed movement to be remotely controlled using magnetic fields.^12^ Magnetotactic sperm cells (MSCs) were obtained using composite magnetic polystyrene microparticles, which selectively bind to the head of the sperm cell, resulting in an unobstructed flagellar beat and a significantly increased swimming speed when compared to earlier sperm-hybrid microrobots.^9,13,25^ Clusters of MSCs furthermore could be detected using ultrasound (US) and photoacoustic (PA) imaging in ex vivo tissues, with microparticles serving as contrast agents, highlighting the potential for tracking these microrobots under in vivo conditions.

In the present study, we optimized the preparation of high-yield, high-quality MSCs and assessed the fertilization potential of bovine sperm bound to magnetic microparticles using conventional IVF. We have improved the purification method (**Figure 1A**) to obtain a larger number of MSCs while separating them from remaining unbound microparticles and particle-free sperm cells. Furthermore, we conducted an extensive assessment of how bound magnetic particles affect sperm functionality, followed by complete IVF cycle experiments with embryo culture up to the blastocyst stage (**Figure 1B**) to demonstrate the feasibility of fertilization using MSCs and to determine possible differences in embryonic development. Additionally, we developed a mechanism for the automated control of MSCs and proposed several strategies to enhance the efficient transport of multiple sperm to the oocyte, including automated image-based sperm recognition and control, sequential transport, and pre-clustering of MSCs before directing them to their target location (**Figure 1C**).

**Figure 1.**
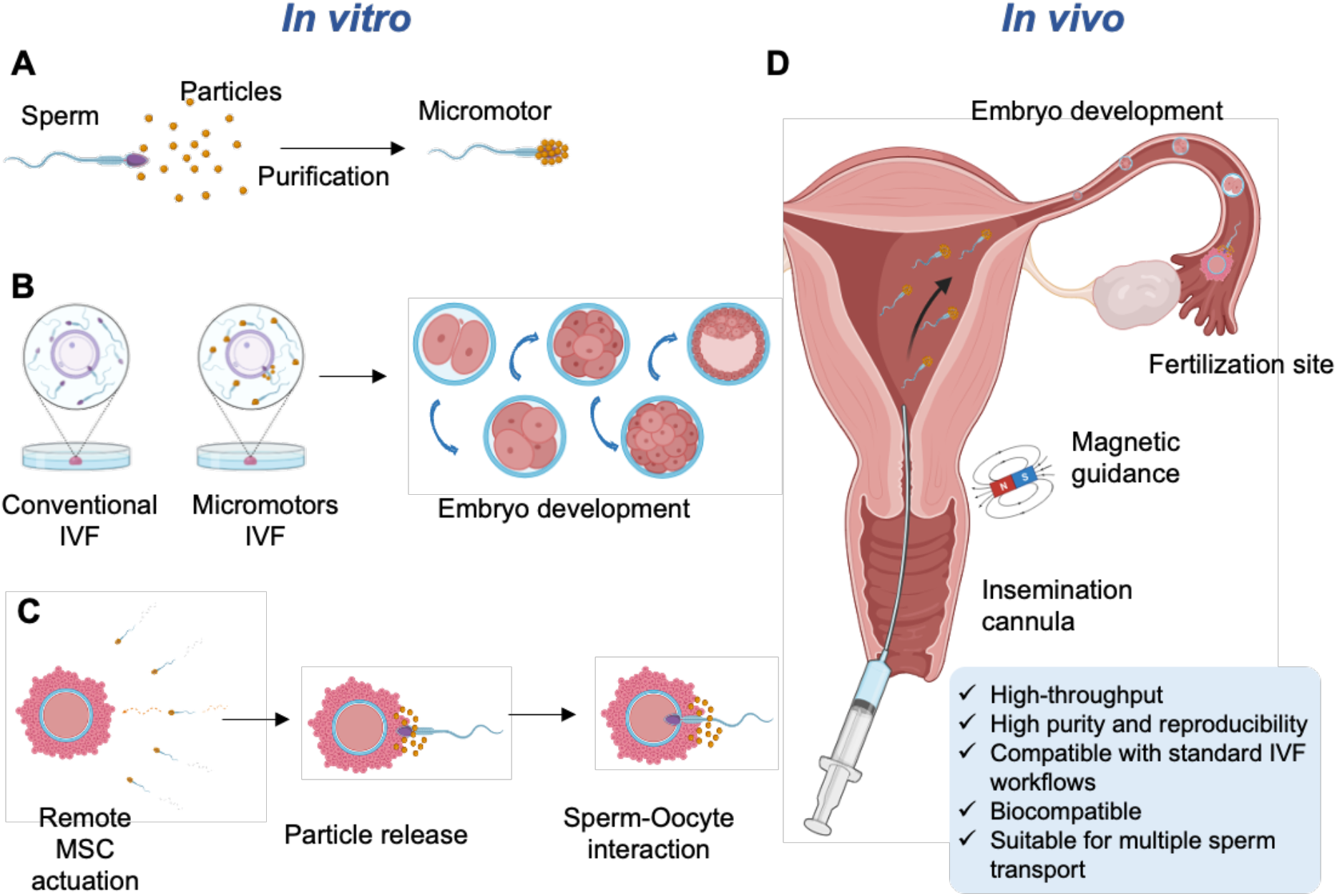
Magnetotactic sperm (MSCs) assembly concept for in vitro fertilization and in vivo assisted insemination. A) Assembly and purification of the microrobots. B) Comparison between conventional IVF and IVF assisted by MSCs, with evaluation of embryo development up to the late-stage blastocyst. C) Remote-controlled navigation of a single MSC toward a cumulus–oocyte complex (COC), followed by particle release during penetration of the cumulus cells, enabling subsequent interaction of free sperm with the oocyte. D) Conceptual illustration for the envisioned intrauterine insemination of MSCs and their magnetic guidance to the ampulla of the fallopian tube for oocyte fertilization. Created with BioRender.

The development of a method for magnetically guided sperm delivery into the female reproductive tract (**Figure 1D**), combined with a suitable imaging method, could significantly reduce the invasiveness of assisted reproductive procedures for women in cases of male infertility. Furthermore, it mitigates the potential and controversially discussed risks associated with conventional IVF procedures^17–19^ by preserving conception within its natural and dynamic microenvironment of the female reproductive tract. This application can be extended to animal species of economic interest or endangered animals for which a functional in vitro fertilization (IVF) is not established.^20,21^

## Methods

### Preparation of sperm samples

Sperm Tyrode’s albumin lactate pyruvate (SP-TALP) medium without Bovine Serum Albumin (BSA; Sigma-Aldrich, St. Louis, MO, USA; Cat. No. A7030) was prepared by adding 2.5 mL of sodium pyruvate (Gibco, Thermo Fisher Scientific, Waltham, MA, USA; Cat. No. 11360070) and 100 µL of Gentamicin sulfate (Sigma-Aldrich, St. Louis, MO, USA; Cat. No. G3632) to 47.5 mL of SP-TL (Caisson Labs, Smithfield, UT, USA; Cat. No. IVL03), the solution was sterile filtered using a 0.20 µm syringe filter (VWR, Radnor, PA, USA; Cat. No. 514-0061) and stored at 4 °C and used within 2 weeks. To prepare SP-TALP with BSA, 300 mg of BSA were dissolved before filtration and storage. Cryopreserved bovine sperm samples from the same individual were obtained from Masterrind GmbH (Meißen, Germany). Sperm straws were thawed in a water bath at 37 °C for 3 minutes, 2 straws were mixed with 2 mL of SP-TALP. 15 mL conical centrifuge tubes (Greiner Bio-One, Kremsmünster, Austria; Cat. No. 188261-N) with a discontinuous density gradient (BoviPure and BoviDilute, Nidacon International AB, Mölndal, Sweden; Cat. No. BP-100/BD-100) of 1 mL 80% BoviPure, 1 mL 40% Bovipure and 2 mL of sperm sample were centrifuged for 12 minutes at 300 g and 37 °C, then the supernatant was discarded, and the pellet was mixed with 3 mL of SP-TALP in a new 15 mL conical centrifuge tube. After centrifugation of 5 minutes at 300 g and 37 °C, pellets were resuspended with sperm media.

### Assembly of MSCs in Petri dishes

Magnetic polystyrene composite microparticles (Sigma-Aldrich, St. Louis, MO, USA; Cat. No.75597) were washed twice and stored overnight with Dulbecco’s Phosphate Buffered Saline (PBS) and antibiotics: Amphotericin B (Pan Biotech, Aidenbach, Germany; Cat. No. P06-01050), Penicillin/Streptomycin (Gibco, Thermo Fisher Scientific, Waltham, MA, USA; Cat. No.15140122) and Gentamicin sulfate, then they were centrifuged and resuspended in ultra-pure water before use. 35 mm Petri dishes (Greiner Bio-One, Kremsmünster, Austria; Cat. No. 627160) were filled with polydimethylsiloxane (PDMS; SYLGARD 184; Farnell GmbH, Frankfurt am Main, Germany; Cat. No. 101697) and cured overnight at 65 °C on a tilted hotplate (VWR, Radnor, PA, USA; Cat. No. 12365-486) elevated in the back 3 mm. Dishes were washed twice with PBS (BioWest, Nuaillé, France; Cat. No. L0615) and antibiotics and left to dry inside a biosafety cabinet overnight before use. 3 mL of SP-TALP without BSA were added to the dish, followed by 10 µL of microparticles and 50 µL of sperm cells in the deepest part of the dish, before incubation at 37 °C and 6% CO_2_ for 30 minutes.

### Magnetic purification of MSCs

After 30 minutes of incubation at 37 °C and 6% CO_2_, a Neodymium disc magnet (N52, 20 mm diameter x 5 mm thickness; Magnet-Kauf, Germany) was placed under the dish to fix the MSCs, and 2 mL of medium (containing unfunctionalized sperm cells) were removed and replaced by 1 mL of fresh SP-TALP with BSA. After removal of the magnet, 200 µL of sample collected from the Petri dish were layered on 500 µL of BoviPure in 1.5 mL centrifuge tubes. The tubes were incubated for 20 minutes at 37 °C and 6% CO_2_, in which the functionalized sperm sediment while non-functionalized sperm and free particles stay in the overlay. Subsequently, then the supernatant/overlay was discarded and the pellet containing the functionalized sperm was resuspended in a new centrifuge tube with 200 µL of SP-TALP with BSA and incubated for another 20 minutes in which the functionalized sperm will sediment. The pellet was collected and resuspended in 30 µL of BO-IVF medium (IVF Bioscience, Bickland Industrial Estate, Falmouth, Cornwall, United Kingdom; Cat. No. 71004).

For IVF, one sample of MSCs was prepared for every drop containing cumulus oocyte complexes (COCs).

### Assessment of free sperm cell concentration

The number of free sperm cells in the microrobot sample was determined in the final sample after purification. 30 µL of MSCs sample were added to a 5 mm circular reservoir of PDMS and heated until immobilization of all sperm cells was reached. Next, the whole sample was imaged and free sperm cells and MSCs were counted. The same was done with the previously reported purification method^12^ to compare it with the magnetic purification method. Additionally, we compared the reproducibility of sperm-particle binding between two batches (B1 and B2) of magnetic PS microparticles for both separation methods. For each condition, three independent replicates (different straws from the same bull) were performed and all MSCs were counted.

### Acrosome staining

Sperm acrosome integrity was evaluated following the protocol by Yániz et al^26^. Briefly, sperm smears were allowed to air dry, then they were fixed with 96% ethanol for 15 minutes and washed with distilled water. For staining, 20 µL of the staining mix, 5 mg/mL Propidium iodide (PI, Sigma-Aldrich, St. Louis, MO, USA; Cat. No. P4864) and 50 mg/mL *Pisum sativum* agglutinin (PSA, Sigma-Aldrich, St. Louis, MO, USA; Cat. No. 0770) in a PBS-based solution, was added to the glass slide (VWR, Radnor, PA, USA, Cat. No. 631-1553), covered with a glass cover slide, and incubated for 20 minutes at room temperature in the dark. Images were taken using a fluorescence microscope (AxioObserver, Zeiss, Oberkochen, Germany) with an excitation wavelength max. of 498 nm and emission wavelength max. of 518 nm (green) and an excitation wavelength max. of 493 nm and detection wavelength max. of 636 nm (red). Three independent replicates were conducted, and a minimum of 230 cells were counted for each replicate.

### Mitochondria membrane integrity staining

Sperm cells were stained using the BioTracker 488 Green Mitochondria Dye kit (Sigma-Aldrich, St. Louis, MO, USA; Cat. No. SCT136) to detect cell viability, metabolic activity and overall cell health. We followed the protocol according to the manufacturer’s instructions with some modifications. 100 µL of sperm sample were mixed with 1µL of staining solution (prepared according to the kit) and 1 µL of PI was added. Samples were incubated for 15 minutes at 37 °C and analyzed using fluorescence microscopy. Pictures were taken using an absorbance wavelength of 490 nm and an emission wavelength of 523 nm (green). The excitation wavelength was maximized at 493 nm, and the detection wavelength was maximized at 636 nm (red). Afterward, sperms were counted. Three independent replicates were conducted, and a minimum of 116 cells were counted for each replicate.

### ROS staining

To evaluate oxidative stress in sperm cells and MSCs, we used the kit CellROX^®^ Green (Invitrogen/Thermo Fisher Scientific, Waltham, MA, USA, Cat. No. C10444). 100 µL of sperm samples were incubated with 2 µL of 5 µM CellROX^®^ reagent for 20 minutes at 37 °C. Samples were centrifuged for 5 minutes at 300 g, and fresh SP-TALP was added. For a negative control, the CellROX^®^ reagent was omitted. For positive control, we induced the ROS production by as described in Celeghini et al.^27^. Briefly, 25 µL iron sulfate (4mM, Sigma-Aldrich, St. Louis, MO, USA; Cat. No. F8633), 25 µL ascorbic acid (20mM, Sigma-Aldrich, St. Louis, MO, USA; Cat. No. A7506), and 50 µL hydrogen peroxide (30%, Sigma-Aldrich, St. Louis, MO, USA; Cat. No. 216763) were added to 100 µL of sperm solution. The sample was incubated with the lid open for 10 minutes at 37 °C before staining. Stressed sperm cells show green fluorescence, an excitation wavelength of 450 to 500 nm, and a detection wavelength of 515 to 565 nm was used for detection. Three independent replicates were conducted, and a minimum of 239 cells were counted for each replicate.

### DNA fragmentation staining

Sperm DNA fragmentation was evaluated using the terminal deoxynucleotidyl transferase dUTP nick end labelling (TUNEL) assay according to the manufacturer’s instructions (*In Situ* Cell Death Detection Kit, Fluorescein; Roche Diagnostics, Mannhein, Germany; Cat. No. 11684795910) with modifications. Sperm smears were prepared, left to air dry, and fixed for 1 hour with the fixation solution. Slides were rinsed with PBS and incubated with the permeabilization solution for 30 minutes at 37 °C and washed again. For the positive control, the slide was incubated with 2385U/mL DNase I for 30 minutes at 37 °C to induce double stranded DNA breaks. For negative control, 50 µL of label solution were added. For the positive control and samples, 50 µL of the TUNEL mix was added to each slide. Slides were covered with Parafilm (Sigma-Aldrich, St. Louis, MO, USA; Cat. No. P7543) to ensure homogeneous spread and incubated for 90 minutes in a humidified chamber in the dark at 37 °C. Samples were analyzed using a fluorescence microscope using an excitation wavelength of 450 to 500 nm and detection wavelength of 515 to 565 nm (green). Fragmented cells appeared green. Three independent replicates were conducted, and a minimum of 74 cells were counted for each replicate.

### Bovine oviduct epithelial cells (BOECs)

Bovine oviducts were obtained from a slaughterhouse (Vorwerk Podemus e.K., Dresden, Germany), and bovine oviductal epithelial cells (BOECs) were isolated and cultured following the protocol described by Palma-Vera et al^28^. In brief, surrounding tissue and blood vessels were removed from the oviducts, and were rinsed with PBS. One end of the oviduct was closed with a clamp, and the lumen was then filled with collagenase 1A (1 mg/mL, Sigma-Aldrich, St. Louis, MO, USA; Cat. No. C2674) prepared in Ham’s F12 medium (Thermo Fisher Scientific, Waltham, MA, USA, Cat. No. 31765027). The other end was closed with a second clamp, and the oviduct was incubated at 37°C for 60 minutes for enzyme digestion in Ham’s F12 medium with antibiotics. One end was cut, and the mucosa was gently squeezed out and passed through a 40 μm pore-size strainer. The retained cell clusters were washed with PBS and treated with 0.5% trypsin/EDTA (Invitrogen/Thermo Fisher Scientific, Waltham, MA, USA, Cat. No. 25300-054) at 37°C for 10 minutes. The reaction was stopped by adding fetal calf serum, and the solution was passed through a strainer once more. The collected cells were either seeded or frozen for future use. 200,000 cells were seeded into 0.4 μm pore-size inserts pre-coated with rat tail collagen I (SERVA Electrophoresis GmbH, Heidelberg, Germany, cat. no. 47254.01) for cell culture. The cells were initially maintained under liquid-liquid conditions in proliferation media for one week to allow cell division. Subsequently, the medium from the insert was removed to establish an air-liquid interface, promoting cell differentiation. The cultures were maintained for at least four weeks before experimental use, with media exchanges performed three times per week.

Free sperm cells or MSCs were added to an insert of ciliated BOECs. Each sample was incubated at 37°C and 6% CO2 for 30 minutes before preparing for SEM imaging.

### Oocyte collection and maturation

Bovine ovaries were collected from a slaughterhouse and transported to the lab in a transport solution kept at 30 °C. Transport solution was prepared by dissolving 9 g sodium chloride (NaCl, Sigma-Aldrich, St. Louis, MO, USA; Cat. No. S5886), 0.66 g Penicillin G (Sigma-Aldrich, St. Louis, MO, USA; Cat. No. P3032), and 0.131 mg Streptomycin sulfate (Sigma-Aldrich, St. Louis, MO, USA; Cat. No. S9137) in 1 L of sterile diH_2_O. Follicles were sliced and follicular fluid was collected with OPU medium (IVF Bioscience, Bickland Industrial Estate, Falmouth, Cornwall, United Kingdom; Cat. No. 51001) in 50 mL tubes. COCs were left to sediment for 10 minutes and the supernatant was discarded, a second wash with OPU medium was performed to remove excessive blood. After the second wash, the sediment was transferred to 100 mm Petri dishes for the selection of COCs. COCs were placed in another 100 mm Petri dish with 10mL OPU medium. Collection was performed in a maximum of 60 minutes. COCs were washed twice with Wash medium (IVF Bioscience, Bickland Industrial Estate, Falmouth, Cornwall, United Kingdom; Cat. No. 51002) and one time with BO-IVM medium (IVF Bioscience, Bickland Industrial Estate, Falmouth, Cornwall, United Kingdom; Cat. No. 71001). Finally, 45 COCs were added to 500 µL of BO-IVM medium in each well of a 4 well plate and incubated for 24 h at 38.8 °C, 6% CO_2_ for maturation.

### IVF

COCs were washed after 24 h of maturation with BO-IVF media and 5-8 COCs were transferred to drops of 50 µL of BO-IVF medium under oil (IVF Bioscience, Bickland Industrial Estate, Falmouth, Cornwall, United Kingdom; Cat. No. 51003) and distributed randomly between six groups. For the Parthenogenetic group (P), only COCs were added, in the group Parthenogenetic with Particles (PP), 1 µL of a 1:10 dilution of magnetic microparticles suspended in BO-IVF media was added to each drop. An IVF control was performed by adding a sperm sample with a final concentration of 2×10^6^ mL^-1^ used as a quality control. The sperm concentration was determined using a Neubauer chamber; if needed, dilutions were done to adjust the required sperm concentration. The magnetotactic sperm cell group (MSC) had a final concentration of 275,000 sperm mL^-1^ after purification and 13,750 MSCs were added to each drop. Medium concentration (MC) and Low concentration (LC) groups underwent the same preparation as the MSCs but without the addition of magnetic particles. The sperm concentration in the MC group was adjusted to the same number as the MSC group and the concentration in the LC group was as adjusted to the number of particle-free sperm cells after purification. After coincubation for 16 h, oocytes were removed from the drop using a pipette (internal diameter = 275 µm, Gynemed GmbH & Co. KG, Oldenburg, Germany, code GY-275/20) and placed in a 35 mm Petri dish with 3 mL of Wash medium for denudation using a smaller pipette tip (internal diameter = 130 µm, Gynemed GmbH & Co. KG, Oldenburg, Germany, code GY-130/20). Oocytes were washed in a drop of BO-IVC medium (IVF Bioscience, Bickland Industrial Estate, Falmouth, Cornwall, United Kingdom; Cat. No. 71005) and then placed in a 50 µL drop BO-IVC medium under oil at 38.8 °C, 6% CO_2_ and 6% O_2_. After 24 hours, cleavage embryos were transferred to a new dish with a 50 µL drop of BO-IVC medium under oil. On day 7, blastocyst development was observed and on day 9 hatched blastocysts were counted. Around 200-300 COCs were used per IVF replicate, and a minimum of 21 oocytes was used per study group.

### SEM imaging

BOECs with sperm cells or MSC samples were washed with Hank’s Balance Salt Solution and fixed in 3% glutaraldehyde in SEM buffer (0.1 M sodium phosphate buffer, pH 7.4 with 0.1 Sucrose) overnight at 4 °C. Samples were washed two times with SEM buffer and then dehydrated in increasing ethanol (Th. Geyer GmbH & Co. KG, Höchstadt, Germany, cat. no. 11991521) concentrations: 35%, 50%, 70%, 95%, and 100%. Samples were then washed with hexamethyldisilazane (Sigma-Aldrich, St. Louis, MO, USA; Cat. No.440191) and left to dry overnight. Samples were coated with a 10 nm layer of gold by physical vapor deposition. Images were taken using a Zeiss DSM 982 microscope with accelerating voltage of 5kV. Cells were counted in all images taken.

### Closed-loop control of magnetotactic sperm cells

Remote guidance of MSCs was achieved through microscopic imaging, real-time tracking and magnetic actuation of the cells in a multimodal closed-loop control system^29^. A digital camera (a2A2590-60ucPRO, Basler AG, Germany) on an inverted microscope (Eclipse Ti2-E, Nikon Corp., Japan) captured monochromatic bright-field images (2592 x 1944 pixels, 8 bits per pixel) at 30 Hz. Real-time image processing for binarization with a configurable threshold and calculation of the centroid was implemented based on CuPy^30^ and performed on a GPU (GeForce RTX 3090, Nvidia, USA). The localized microrobots were linked between consecutive frames using TrackPy^31^ running on CPUs (2x Intel Xeon Gold 5217, Intel, USA). One MSC was selected for closed-loop control, for which the difference vector to the target location was calculated continuously and used as direction of the magnetic field vector. The magnetic field with a strength of 10 mT was generated with a commercial 8-coil setup (MFG-100-i, Magnebotix AG, Switzerland).

### Statistical Analysis

If not indicated otherwise, error bars represent the standard deviation. In box plots, median values are represented by a line and mean values by a circle and were done with Origin (OriginLab Corporation, Northampton, MA, USA). The statistical analysis for mean, standard deviation and t-test was performed using Microsoft Excel (Microsoft Corporation, Redmond, WA, USA). For remote control experiments, we used an unpaired two-sample t-test, implemented in SciPy^32^. Differences were considered significant when p < 0.05.

## Results and Discussion

### Optimization of MSC purification

In our previous work, we developed an assembly and purification method for MSCs used in IVF. We will refer to this method as the basic separation method (BS) in this manuscript **Figure 2A**). However, the final samples still contained unbound spermatozoa^12^. To further reduce the number of particle-free sperm cells in the sample, we improved the BS and named it the magnetic separation method (MS; **Figure 2A**). A thin layer of PDMS was added to the bottom of the petri dish and cured in a tilted position. Then, a magnet was placed under the petri dish after the assembly of MSCs, and the sperm medium was exchanged magnetically separating particle-bound and particle-free sperm cells. The PDMS layer served two functions: to accumulate microparticles in the deepest part of the petri dish, where the spermatozoa will be placed to facilitate the contact between them, and to serve as a spacer between the magnet and the sample to prevent damage to the spermatozoa. Additionally, we compared the reproducibility of sperm-particle binding between two batches (B1 and B2) of magnetic PS microparticles (**Table S1 and S2, Supporting Information**).

**Figure 2.**
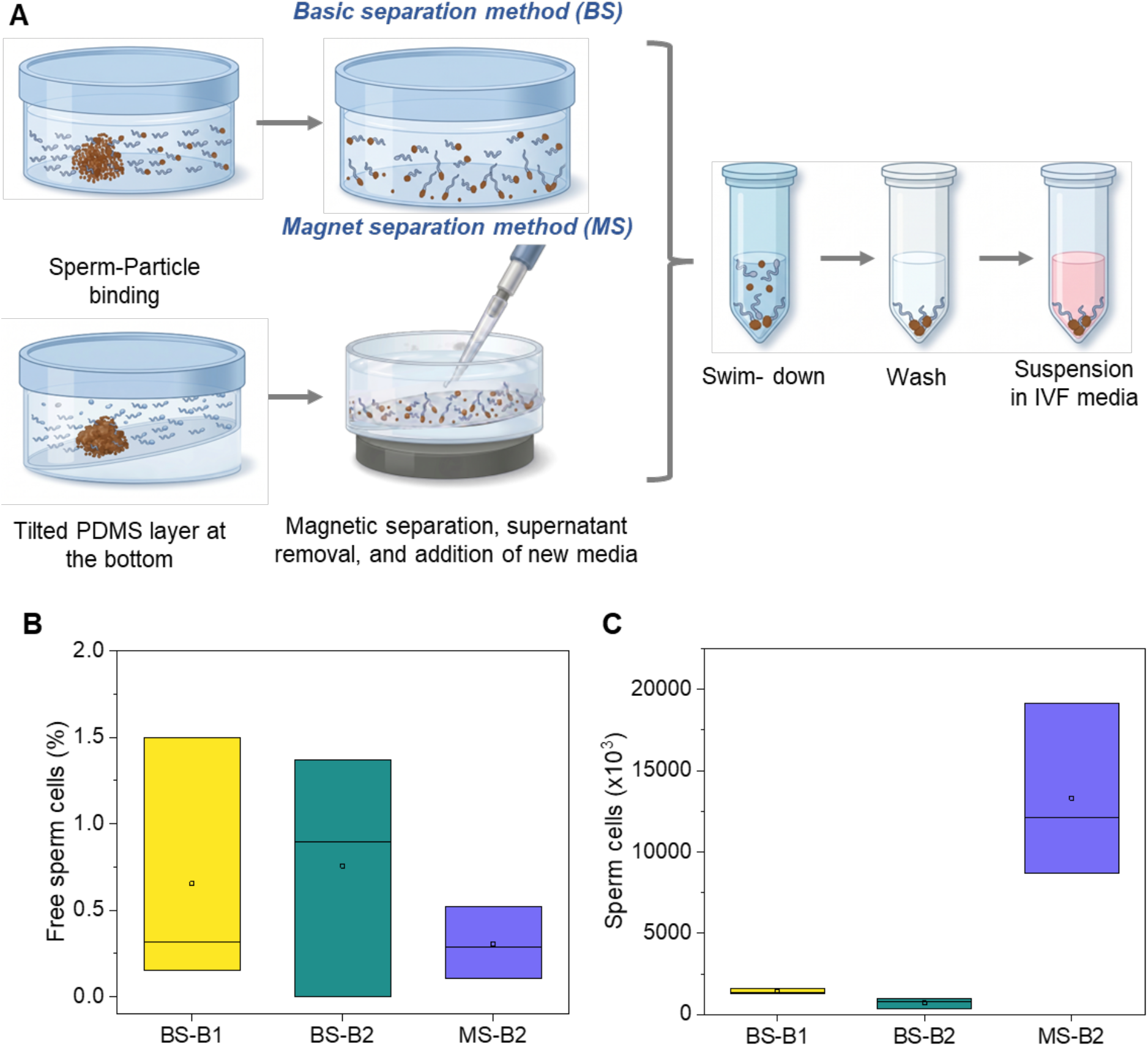
Optimized method for obtaining highly pure MSCs for IVF and the comparison with the previously reported protocol^12^, Basic separation method (BS). A) Two methods for the purification of MSC: BS and Magnetic separation (MS). B) Percentage of particle-free sperm cells after purification using the BS and MS techniques with two batches of magnetic microparticles (B1 and B2). C) Final number of MSCs obtained after purification using the BS and MS techniques with two batches of magnetic microparticles (B1 and B2).

We did not observe any significant differences in the number of free sperm between the two batches of particles using the basic separation method (B1: mean 0.65 ± 0.73% SD, B2: mean 0.75 ± 0.69% SD) (**Figure 2B**). MS reduced the fraction of particle-free sperm to 0.30 ± 0.20%. Crucially, MS yielded substantially more MSCs (13,267 ± 5,332) than the BS methods (BS-B1: 1,404.33 ± 153; BS-B2: 709.33 ± 334.74) (**Figure 2C, Table S3, Supporting Information**). This roughly tenfold increase in MSC yield is critical for IVF experiments, which typically require millions of sperm for preparation and sperm:oocyte ratios >10,000:1 in dish-based fertilization^28^.

### Effect of particle-functionalization on sperm health

To assess the effect of the binding of microparticles on sperm health, we performed different assays to monitor acrosome integrity, mitochondria membrane integrity, oxidative stress and sperm DNA fragmentation at 0 h and 2 h of incubation after the assembly of MSCs and compared the results to a control sample of particle-free sperm cells incubated under the same conditions. Even though larger differences between the two samples could be expected at longer time periods, we estimate that possible applications of MSCs in assisted reproduction would not last longer than 2 hours and therefore focus our analysis on this period. This conservative time duration was estimated by considering the swimming speed of MSCs (121.1 ± 11.7 µms^−1 12^) and the length of the oviduct (11-12 cm in humans^33^ and 25 cm in bovine^34^). The estimated travel time for a human oviduct is approximately 17 minutes, while for a bovine oviduct, it is around 34 minutes. Even though the movement of spermatozoa in the reproductive tract might be substantially slower and the effect of magnetic guidance in vivo is yet unknown, the overall procedure workflow is designed to accommodate the estimated time, ensuring efficient and effective treatment.

#### Acrosome integrity

Acrosome integrity was assessed by a fluorescent staining using the lectin *Pisum Sativum* agglutinin (PSA) conjugated with fluorescein isothiocyanate (FITC). PSA has affinity for r α-D-mannose and α-D-glucose and recognizes glycosylated proteins present in the acrosome^35,36^ (**Figure 3A**). The acrosome plays a fundamental role in fertilization as it allows the sperm to recognize the oocyte and contains the enzymes necessary to digest the cumulus oophorous and the zona pellucida before fusion with the vitellus^37^. We classified the sperm cells by degree of staining of the acrosome: intact acrosome, low staining, clearly damaged and totally lost (**Figure S1A, Supporting Information**). The changes in staining brightness are related to alterations in the acrosome, which can be partial or complete due to acrosome reaction. In both groups, control and MSC, we observed a reduction of the number of sperm cells with intact acrosome after 2h of incubation, from 70.31 ± 9.88% and 78.73 ± 1% to 53.98 ± 4.06% and 56.19 ± 5.83%, respectively (**Table S4, Supporting Information**). No significant differences were observed between both groups at 0 h and 2 h, indicating that acrosome integrity is not negatively affected by binding of magnetic particles.

**Figure 3.**
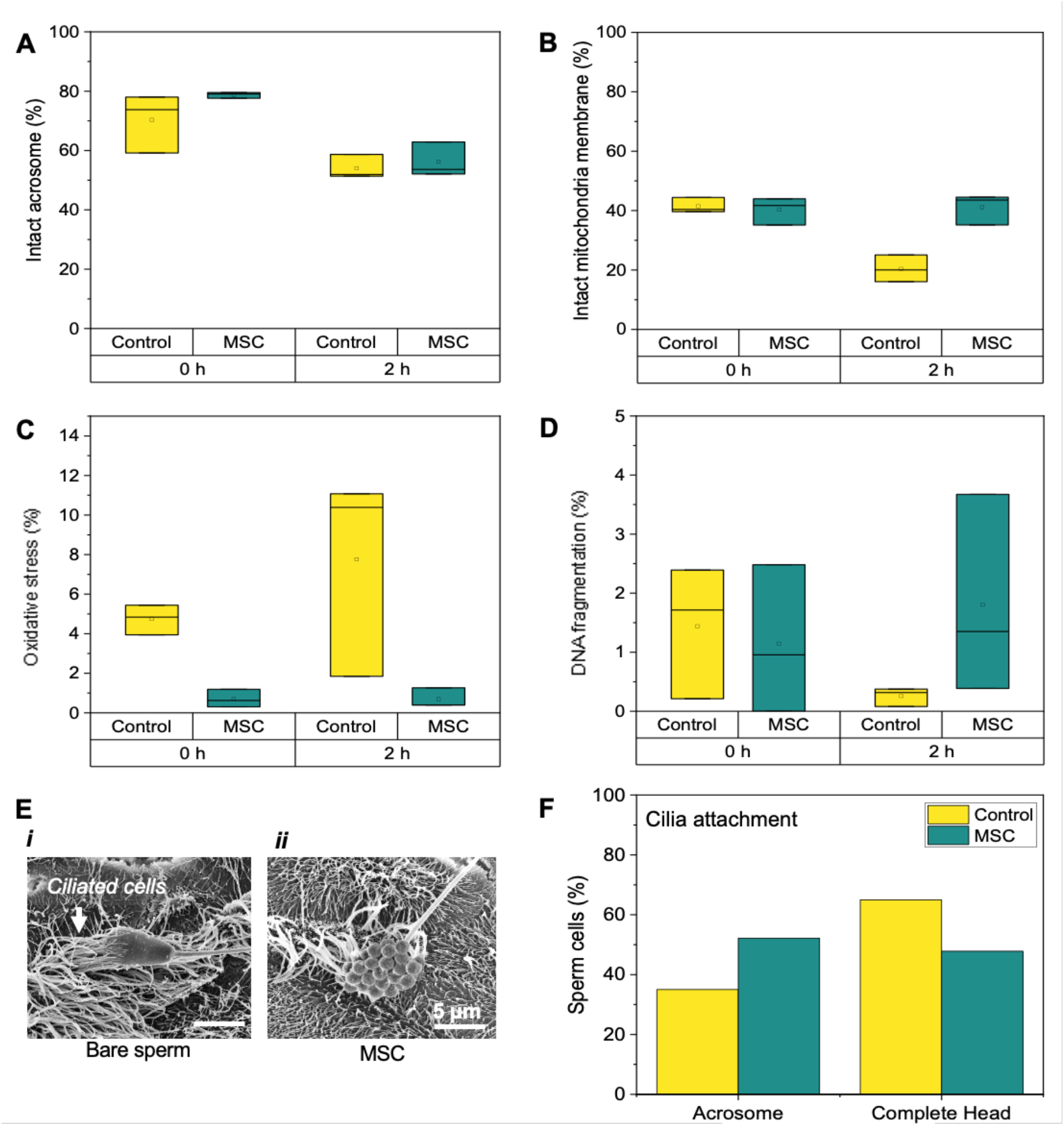
Effect of microparticle-binding on sperm health. A) Acrosome integrity of MSCs compared to particle-free sperm cells at 0 h and 2 h of incubation, measured by fluorescently labelled PSA and PI. B) Mitochondria membrane integrity of MSCs compared to particle-free sperm cells at 0 h and 2 h of incubation, measured by BioTracker 488 Green Mitochondria Dye. C) Oxidative stress of MSCs compared to particle-free sperm cells at 0 h and 2 h of incubation, measured by using CellROX™. D) DNA fragmentation of MSCs compared to particle-free sperm cells at 0 h and 2 h of incubation, measured by using TUNEL assay. E) Representative SEM images of the attachment of BOEC cilia to the sperm head of particle-free sperm cells (i) and MSCs (ii). F) Location of cilia attachment on the sperm head in MSCs and particle-free sperm cells.

#### Mitochondria membrane integrity

Mitochondria membrane integrity was also evaluated using a fluorescent staining (**Figure 3B**). The membrane-permeable dye accumulates in the membrane of mitochondria in metabolically active and living cells, but the signal decreases as the mitochondria membrane disintegrates during cell death. Additional staining with PI allows for evaluation of cell health metabolic activity and cell viability. Sperm cells were classified into three groups (**Figure S1B, Supporting Information**): intact membrane (metabolically active and live cells), defective membrane (low metabolic activity and dying cells) and damaged membrane (no active metabolism and dead cell). We observed a reduction of the portion of cells with intact mitochondrial membrane in the control sample after two hours of incubation, from 41.48 ± 2.57 % to 20.38 ± 4.50 % (**Table S5, Supporting Information**). In contrast, mitochondria membrane integrity remained stable in the MSC group (40.27 ± 4.57 % to 41.07 ± 5.12 %) after two hours of incubation and was significantly higher at 2 h than in the control sample. In the case of the other two sperm classifications: defective membrane and damaged membrane we found significant differences between the control and MSC. For the defective membrane group, the control group had a lower percentage of sperm with this defect after 2 hours of incubation (17.98 ± 12.53%) compared to the MSC group (52.21 ± 0.83%), which was statistically significant (*p* = 0.009). In the damaged membrane group, the control group was higher than the MSC after 2h of incubation, 61.63 ± 17.04% vs 6.7 ± 4.45% (*p* = 0.005). These results point to a delayed cell death in the MSC group, which is consistent with previous results where cell viability in the MSC group after 6 hours of incubation was significantly higher than in the control group^12^. These results could be explained by an absorption of BSA present in SP-TALP (sperm media) on the polystyrene microparticles after the sperm-microparticle binding. The binding of BSA will form a protein corona around the polystyrene microparticle, as previously reported by Kihara et al. ^38^. Albumin has antioxidant properties^39^ and is known to be preferably absorbed onto hydrophobic surfaces^40^, like the silane layer of the magnetic particle, thus potentially reducing the free radicals around the sperm cell.

### Oxidative stress

To further understand the positive effect of the PS microparticles on sperm viability and mitochondria membrane integrity, we evaluated the oxidative stress in both groups. The used dye, CellROX™, accumulates inside the cell; in a reduced state the fluorescence is weak and exhibits bright green photostable fluorescence upon oxidation by reactive oxygen species and subsequent binding to DNA (**Figure S1C, Supporting Information**). It is estimated that 90% of reactive oxygen species can be traced back to the mitochondria^41^. The initial values at 0h of oxidative stress in the control group were higher than in the MSC group, 4.98 ± 0.82 % vs 0.70 ± 0.44 %, respectively (**Figure 3C** and **Table S6, Supporting Information**). After 2h of incubation, the control group had a higher percentage of oxidative stress-positive sperm cells (7.76 ± 5.14 %) than the MSC group (0.68 ± 0.50 %), but the difference was not significant between both groups. It is important to note that, compared to the initial values, oxidative stress in the MSC group remained almost unchanged, while in the control group, the values almost doubled. The results in the control group after 2h of incubation are consistent with the percentage of intact mitochondria membrane, meaning that damage to mitochondria membrane is correlated to higher levels of oxidative stress.

#### DNA fragmentation

Sperm DNA fragmentation was determined using a direct and sensitive method called TdT-mediated dUTP-X nick end labeling (TUNEL) where single and double stranded DNA breaks are labeled. The single or double stranded DNA breaks are detected by labeling the free 3′-OH terminal with modified nucleotides, fluorescein-dUTP, in an enzymatic reaction^42^. Non-fragmented sperm DNA will not show any fluorescence, whereas DNA-fragmented sperm cells will show a green fluorescence in their head (**Figure S1D, Supporting Information**). We did not observe any significant differences in the portion of sperm cells with fragmented DNA between the control sample and the MSC sample at 0 h (Control: 1.43 ± 1.11%, MSC: 1.14 ± 1.25%) or after 2 h of incubation (Control: 0.25 ± 0.15%, MSC: 1.80 ± 1.68%) (**Figure 3D** and **Table S7, Supporting Information**). We observed remarkably low levels of sperm DNA fragmentation across both groups, these findings suggest that short-term exposure to the used magnetic microparticles does not compromise sperm DNA integrity, thereby supporting the safety of this co-culture approach. Interestingly, the control group exhibited a lower mean percentage of fragmented sperm at 2 h compared to 0 h. Considering the very low absolute values, where small counting differences have a high relative impact, this reduction is most likely attributable to assay variability.

#### Oviduct epithelium binding

To gain a better understanding of the movement of MSCs in a relevant environment, we studied the interactions between MSCs and bovine oviduct epithelial cells (BOECs), focusing on possible interferences with the sperm-cilia binding by the presence of magnetic microparticles. It has been shown that sperm cells are able to maintain their motility and fertilizing capability in the oviduct for several days by creating a sperm reservoir by binding between the acrosome and the ciliated epithelium^43^. Sperm binding to cilia is mediated by the carbohydrate moieties on the cilia, specifically fucose in bovine^44^. BOECs were extracted from samples collected from a slaughterhouse and frozen until use. During the first week, thawed cells were cultured in inserts within well plates at a liquid-liquid interface to allow cell division and monolayer formation. For the next three weeks, BOECs were cultured at an air-liquid interface to promote the differentiation into ciliated and secretory cells. After differentiation, MSCs or particle-free sperm cells were added in an equivalent amount to the inserts. The samples were then incubated for 30 min before fixation, dehydration and drying. We acquired SEM images of both samples (**Figure 3E i and ii)** to evaluate the interaction between the sperm head and ciliated cells. We observed the attachment of cilia to the sperm head in both samples which could indicate that the presence of magnetic microparticles on the sperm head does not interfere strongly with the cilia trying to adhere to the sperm head, since they try to find their way between the particles to touch the sperm head. It seems like there was no strong preference for cilia adhesion in the acrosomal region specifically or the complete head (**Figure 3F** and **Table S8, Supporting Information**), but further systematic studies are required to elucidate the mechanism of sperm attachment in the MSC sample due to the limited number of studied cells. A weaker binding to the cilia of the oviduct epithelium could be beneficial for the use of MSCs in vivo, as it would not interfere with magnetically guided delivery of sperm cells within the oviduct. Moreover, incomplete coverage of the sperm head with microparticles likely allows the sperm cell to interact with the oviduct fluid, which has a beneficial impact on sperm motility, capacitation, acrosome reaction, and sperm survival^45,46^.

### Use of MSCs in in vitro fertilization

To evaluate the fertilization capacity of sperm cells after binding to the magnetic microparticles, we created different study groups (**Figure 4A**) to perform IVF and subsequently in vitro embryo culture (IVC) until blastocyst stage (**Figure 4B-D**). A Control IVF (C.IVF) served as a control of any effects of media, gamete quality, or other external factors during IVF. A native sperm concentration of 2×10^6^ sperm/mL was used in this group as recommended by the manufacturer of the IVF medium suite (see Materials and Methods section). The MSC sample was prepared and purified as described earlier (**Section 1**), resulting in a final concentration of 275,000 sperm mL^-1^ in the drop of 50 µL. Medium concentration control samples (MC) and low concentration control samples (LC) underwent the same preparation and selection process as the MSC sample but without the addition of any magnetic microparticles. Sperm concentration in the MC group was adjusted to the same concentration as in the MSC group to compare fertilization rates when using MSCs and particle-free cells at the same concentration. In the LC group, the sperm concentration was further decreased to a number similar to the remaining particle-free sperm cells in the MSC sample (∼50 sperm cells) after performing the MSC purification, in order to assess the chances of fertilization by these remaining cells. Higher sperm-oocyte ratios are associated with higher oocyte cleavage and blastocyst rates^47^. Sperm-oocyte ratios used were: 20,000:1 (C.IVF), 2,750:1 (MC and MSC) and 10:1 (LC). Additionally, we prepared two IVF groups without the addition of any sperm cells to monitor parthenogenetic oocyte division. In bovines and other mammals, oocyte cell division can be triggered without prior fertilization by a sperm cell (parthenogenesis), e.g. by physical or chemical stimuli such as mechanical stress, temperature shocks, electrical pulses, ionic treatments, chemical activators, or enzymatic exposure.^48,49^. Oocytes were cultured without any further additions in the parthenogenetic group (P), while 4.3×10^6^ magnetic microparticles were added to the culture in the parthenogenetic + particles (PP) group. P and PP groups served as a control of parthenogenetic oocyte activation and development during IVF in general or stimulated by the presence of the used microparticles. Assuming the size of an oval sperm head, with a length of 9 micrometers and a width of 4 micrometers^50^, we can calculate the area of an ellipse and multiply it by two (the two sides of the sperm head) to estimate the surface area. This calculation allows approximately a maximum of 70 particles to cover the entire head of the sperm. We added approximately 14 times more particles to the PP group, to ensure that we have the same number of particles as in the MSC group, including the particles attached to the sperm cells any unbound particles remaining in the sample after purification.

**Figure 4.**
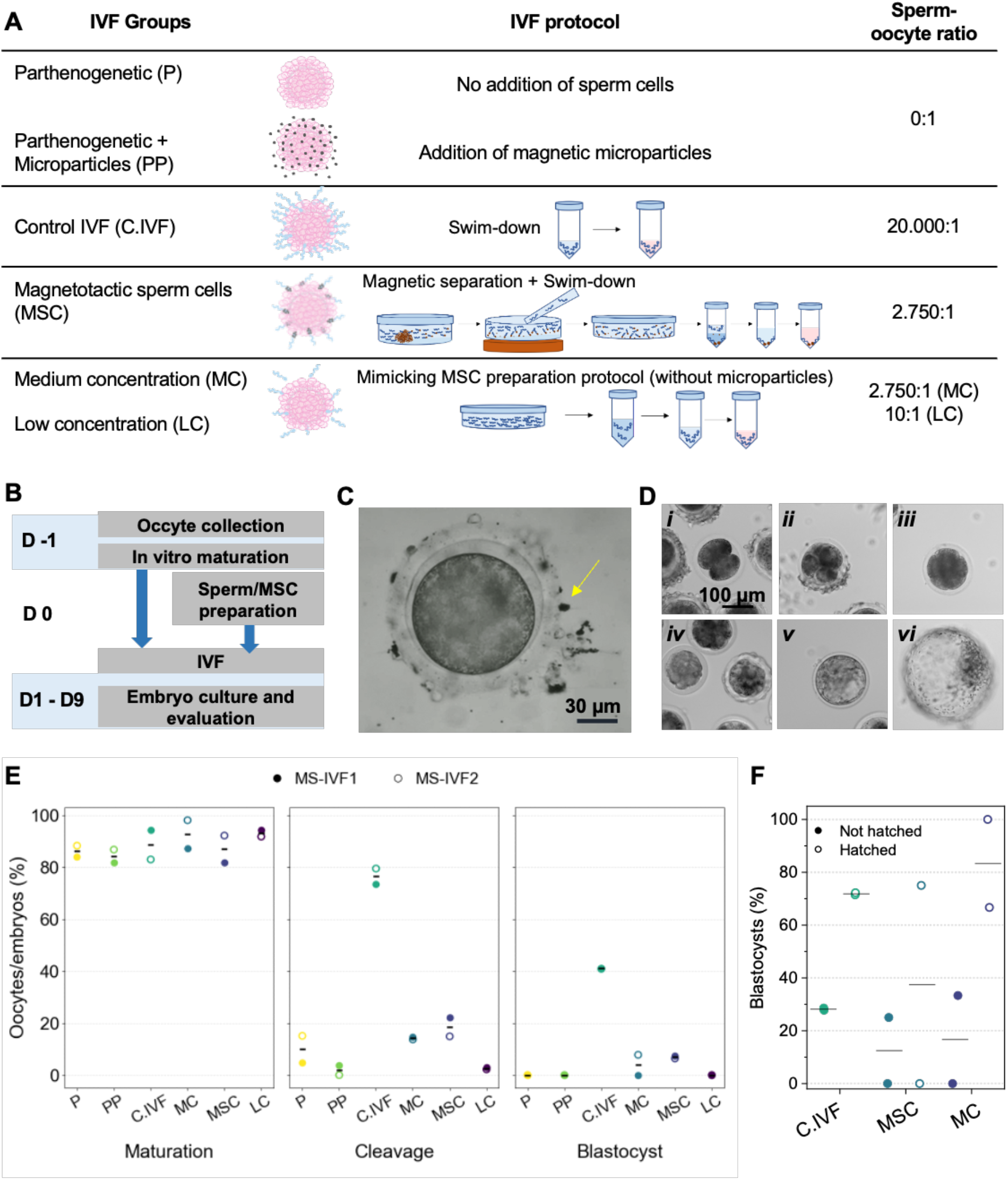
IVF outcomes using MSCs compared with their respective control groups. A) Different IVF groups: control samples and the preparation of MSC, with their specific characteristics and sperm-oocyte ratio. B) Timeline of the IVF workflow. C) Image of a MSC (yellow arrow) attaching to the cumulus layer of a COC. D) Images of embryo development in a MSC sample: two cells embryo (i), four cells (ii), eight cells (iii), morulae (iv), blastocyst (v) and hatched blastocyst (vi). E) Maturation, Cleavage and blastocyst rates observed after using sperm free samples (P and PP), control samples (C.IVF, MC and LC) and MSC. F) Status of blastocysts on day 9 after fertilization to monitor hatching from the zona pellucida.

Ovaries were collected from a nearby slaughterhouse and kept at 30°C until slicing, washing and collection of COCs. After in vitro maturation (IVM) for 24 h, mature oocytes were randomly distributed to the IVF groups. All oocytes with a polar body or divided at day 2 were classified as mature, all empty or atretic oocytes were discarded. Although COCs were randomly allocated to different groups, we found slight differences in maturation rate between groups, with a significant difference between the PP and LC groups (**Figure 4E** and **Table S9, Supporting Information**). This is likely a statistical artifact, and we expect to see similar maturation rates when using larger numbers of COCs.

Fertilization was carried out in drops containing 5-8 COCs and MSCs attached quickly to the cumulus layer. To reduce variability introduced by the sperm sample, all IVFs were performed using samples from the same bull and more specifically from the same lot^51^. We observed the release of microparticles from the sperm head when moving against the cumulus cells and zona pellucida. This likely occurred due to the strong movement generated by the sperm flagellum pushing the cell forward. Additionally, the morphological changes of the sperm head during the acrosomal reaction induced by contact with the cumulus cells could aid in releasing the microparticles^12^, which either attach to the cumulus cells or are released into the media. Oocytes and sperm cells were coincubated for 16 h, before zygotes were denuded and transferred to IVC and embryo development was monitored until hatching of blastocysts.

All presumptive zygotes with two polar bodies or division on day 2 were counted for cleavage rates (Cleavage rate = number of cleavage embryos/number of mature oocytes). As expected, we observed the highest cleavage rate in the C.IVF (76.5 ± 4.25 %) (**Figure 4E**), which is within the range of reported values in bovine IVF^52^. In the MSC group, 18.5 ± 5.1 % of embryos reached cleavage stage, slightly more than in the MC group (14.2 ± 0.6 %), pointing to successful fertilization even after sperm cells were bound to the magnetic microparticles. In contrast, the cleavage rate in the LC group was lower (2.6 ± 0.5 %), likely due to the low number of sperm cells, which indicates that the remaining number of particle-free cells in the MSC sample are unlikely to be responsible for the observed cleavage rate in MSC group. We observed cleavage of embryos in both parthenogenetic groups (P: 9.9 ± 7.3 % and PP: 1.8 ± 2.6 %), but we did not observe any increase of spontaneous oocyte division stimulated by the addition of magnetic microparticles to COCs, as indicated by the low cleavage rate in the PP group.

Embryo culture was performed in 50 µL drops containing all cleavage embryos of a group until blastocyst stage was reached. Parthenogenetic embryos in both groups did not develop into blastocysts but their development was arrested at an earlier stage (**Figure 4E**). Likewise, no embryos in the LC group did develop until blastocyst stage and we cannot exclude that any fertilization observed in the LC group was in fact due to parthenogenetic activation. In the three remaining groups, we observed embryo development up to the blastocyst stage. In line with the results of embryo cleavage, the blastocyst rate (number of blastocysts/number of mature oocytes) was highest in the C.IVF (41.0 ± 0.1 %), followed by the MSC group (6.8 ± 0.7 %) and the MC group (3.9 ± 5.5 %; **Figure 4E**). The similar rates in blastocyst rate for MSC and MC groups suggest a comparable embryo development in both groups, following the similarities in cleavage rates, and the presence of magnetic microparticles used in MSCs does not seem to impede embryo development up to this point. We observed higher rates at all stages of embryo development in the C.IVF group which is even more apparent when comparing embryo development from the cleavage to the blastocyst stage (**Figure S2, Supporting Information**). In the C.IVF group 53.7 ± 3.2% of cleavage embryos developed into blastocysts, followed by the MSC group (38.0 ± 6.7%) and the MC group (28.5 ± 40.4), while in both parthenogenetic groups and the LC group no blastocyst development was observed. To further evaluate embryo development, we monitored the hatching of blastocysts (**Figure 4F**). On day 9 of embryo culture, blastocysts in the Control IVF group (71.8 ± 0.5%), MC group (37.5 ± 53.03 %), and MSC group (83.3 ± 23.57 %) were hatched from the zona pellucida, which does not point to any delayed embryo development in embryos produced using MSCs compared to other groups (**Table S10, Supporting Information**).

Overall, the results of the performed IVF show that bovine sperm cells retain their fertilization capability after the functionalization with magnetic microparticles in the assembly of MSCs. The number of MSCs used for IVF was lower than the number of sperm cells used in the IVFC group. Cleavage and blastocyst rates were comparable in samples produced by MSCs and particle-free sperm cells of the same sperm concentration, whereas the number of remaining particle-free sperm cells in the MSC sample was most likely too small to result in oocyte fertilization (see LC group). Furthermore, any parthenogenetic oocyte division did not lead to development of blastocysts and is therefore unlikely to be the cause of the embryos observed in the MSC group.

While these initial results are promising and demonstrate the effectiveness of MSC in fertilizing an oocyte, one limitation is the relatively low number of replicates (n=2). Despite this, the experiments consistently yield outcomes that support the validity of MSC’s ability to fertilize an oocyte. To strengthen statistical robustness and confirm the reproducibility of these results, increasing the number of replicates will be crucial for future studies.

### Closed-loop control of MSCs and swarm formation

Since a higher sperm-to-oocyte ratio increases the chances of fertilization, we investigated different strategies to control and deliver MSCs to a target: 1) Sequential guidance of single MSCs and 2) Formation and control of MSC clusters. To evaluate the efficiency of closed-loop sperm guidance, a MSC sample was prepared and subjected to alternating intervals of free sperm movement (60 s) and activated guidance (60 s) (**Figure 5A and B**, respectively). The number of MSCs reaching the perimeter of the central target region (defined as a circular area with a 20 µm diameter) was quantified (**Figure 5C**). Under closed-loop guidance, a total of 343 MSCs reached the target region compared to 179 in the control condition, corresponding to a mean of 11.43 versus 5.97 MSCs per interval for 20 intervals each. This represents a 91.6% increase in successful targeting. Statistical analysis confirmed that this increase was highly significant (unpaired two-sample t-test, N=20, t = 5.26, p = 2.2 × 10^−6^; **Figure 5D**).

**Figure 5.**
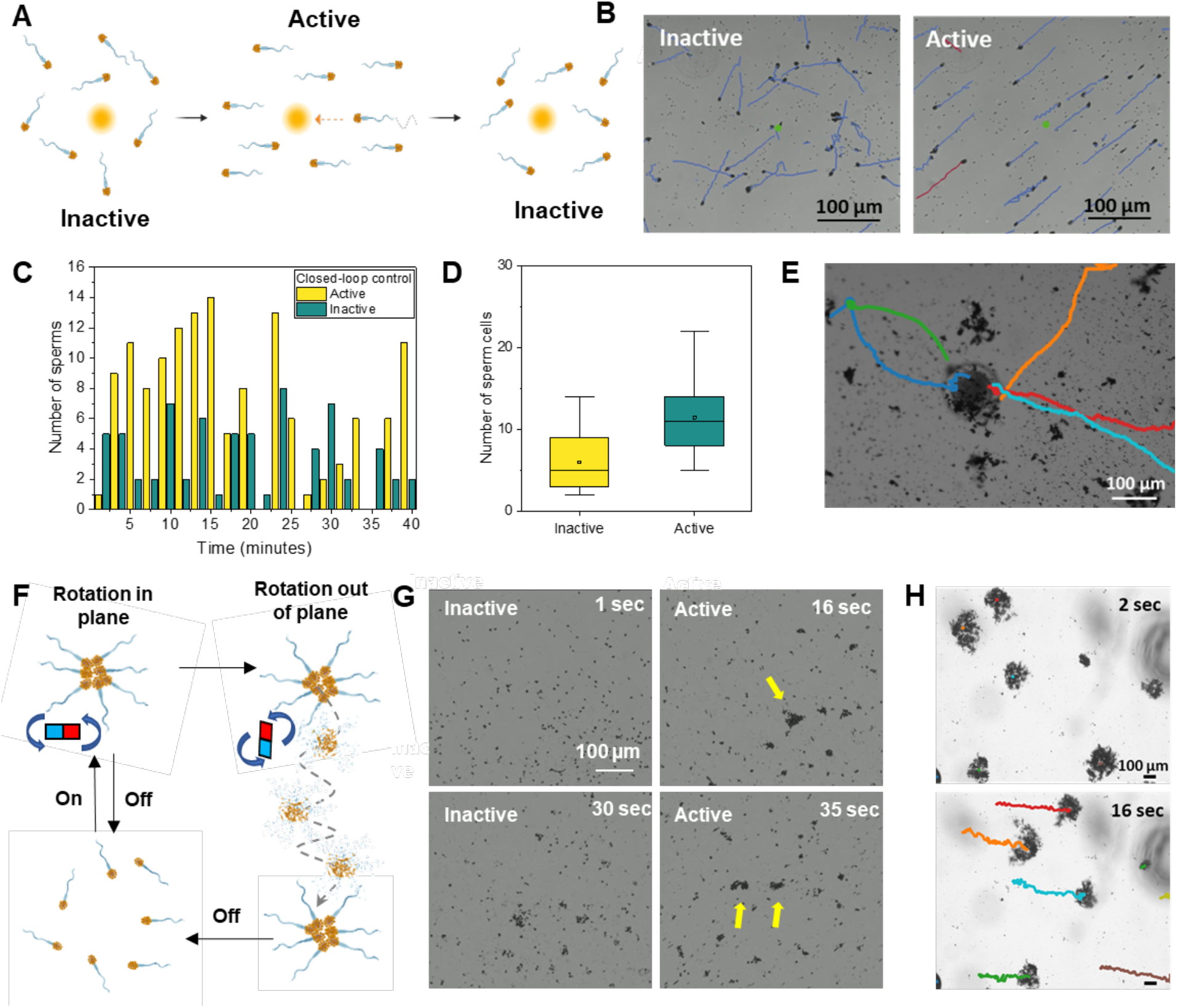
Magnetic control of single MSCs and MSC clusters. A) Schematic of guidance of a specific sperm cell into a target area. B) Images of the trajectories of MSCs while the magnetic control is inactive or active with an MSC selected for guidance in red. C) Number of MSCs guided to the target with closed-loop control alternatingly activated for 1-minute intervals for a total time of 60 minutes. D) Average number of MSCs that reach the target when the Closed-loop control is active and inactive. E) Image of an oocyte with different MSC trajectories to reach the oocyte sequentially. F) Schematic for sperm cluster formation and movement. G) Real images of MSC cluster formation when the magnetic field is active and the disaggregation when it is inactive. H) Multiple MSC clusters with their trajectories over time. Created with BioRender (A and F).

To further demonstrate the guidance towards an oocyte (as a proof of concept), a freshly prepared MSC sample was placed in the closed-loop control system, with a bovine oocyte at the center of the field-of-view. Selected trajectories of MSCs after 110s of closed-loop control toward the oocyte are depicted in **Figure 5E**, showing that five sperms reached the close vicinity in a sequential control scheme.

To increase transported sperm numbers, match the oocyte:sperm ratios used in prior experiments for statistical comparison, and improve visibility for future in vivo guidance, MSCs were aggregated into swarms. **Figure 5F** gives an overview of the magnetic assembly, guidance and disassembly of MSC clusters. A spiral-trajectory magnetic field configuration with B=10 mT was applied with decreasing rotating frequency to 3 Hz over a 300 s interval. Under low-motility conditions, the magnetic attractive forces dominated propulsion, enabling local aggregation into circular cluster patterns. After the 300 s formation phase, the field parameters (10 mT, 3 Hz) were held constant to stabilize the clusters. Microscopic images from the reversible formation process are given in **Figure 5G**. The transport of fully formed clusters over the course of 16s with closed-loop control is shown in **Figure 5H**.

Both sperm guidance strategies are promising but face distinct challenges. Single-sperm guidance demands advanced, potentially parallel control to manipulate individual cells while transporting enough for reliable fertilization. Swarm-based guidance must contend with sperm propulsion opposing magnetic aggregation; effective swarm formation may therefore require transiently suppressing metabolism during long-distance transport and reactivating motility near the oocyte. Pursuing sophisticated swarm formation protocols that preserve sperm viability and function, through controlled metabolic modulation, optimized magnetic protocols, and refined physiological support, offers a particularly attractive route, but will require further investigation and more complex experimental platforms to ensure reliable, fertile outcomes.

## Conclusions

Microrobotic cell/cargo delivery is an active field of research and biohybrid microrobots are becoming more important due to their biocompatibility and unique capabilities. Sperm–hybrid microrobots show promise for assisted reproduction and targeted drug delivery within the reproductive tract. Magnetic composite microparticle–based MSCs enable simple, rapid, and scalable production, making them a practical option for assisted fertilization in vitro and for potential future in vivo implementation. Using a combination of magnetic and density gradient separation, MSCs can be produced and purified in sufficient amounts to ensure oocyte fertilization in vitro. Binding of magnetic microparticles to sperm did not adversely affect the sperm membrane or cellular function. Acrosome integrity, DNA fragmentation, mitochondrial membrane potential, and oxidative stress remained largely unchanged after exposure to the particles; any potential benefit from reduced ROS warrants further investigation. We also examined MSC binding to oviduct epithelium mediated by BOEC cilia and observed no immediate differences compared with particle-free sperm. However, additional studies are required to clarify MSC–oviduct epithelial interactions and to optimize microrobot performance in vivo.

To enhance magnetic control of MSCs for in vitro and in vivo use, we propose several strategies to transport multiple sperm more efficiently toward the oocyte: automated sperm image recognition and guidance, sequential transport, and pre-clustering MSCs prior to relocation. In addition to increasing fertilization probability, employing MSC swarms is expected to markedly improve optical imaging contrast and photoacoustic response.^53^ Together with more advanced planning algorithms^,54^ automated control of MSCs has the potential to be translated to in vivo applications.

To demonstrate the fertilization capability of sperm cells after binding to magnetic microparticles, we carried out IVF using MSCs and the corresponding control groups. The results indicate that embryos could be produced at a reduced rate in the MSC group, even with lower sperm-oocyte ratios compared to conventional IVF. We observed similar cleavage and blastocyst rates in IVF carried out by MSCs and particle-free sperm cells of the same sperm concentration, while low concentration and parthenogenetic control samples allow us to rule out remaining particle-free sperm cells or parthenogenetic activation as the sole cause of fertilization in the MSC group. Embryo development up to blastocyst stage was apparently not impeded in the MSC group, but a deeper analysis of embryo development, ploidy, and possible genetic effects should be investigated in a follow up study. Furthermore, implantation and pregnancy rates of embryos produced in vitro using MSCs should be studied in animal models.

In conclusion, this work demonstrates the feasibility of oocyte fertilization in vitro using MSCs and embryo development up to the blastocyst stage. These results highlight the potential of sperm-hybrid microrobots for assisted reproduction, particularly for precise, guided artificial insemination targeted to the fertilization site in the fallopian tube, which could become an alternative treatment for patients with infertility.

## Acknowledgments

M. Medina-Sánchez thanks the financial support received from the European Union’s Horizon 2020 research and innovation program (ERC Starting Grant Nr. 853609), the HORIZON-MSCA-2022-COFUND-101126600-SmartBRAIN3, and the Spanish Ministry of Science, Innovation, and Universities through the ‘Generación de Conocimiento’ program (Project reference: MICIU/AEI/ 10.13039/501100011033).

We thank Dr. Bergoi Ibarlucea (Tecnalia Research & Innovation, San Sebastián, Spain) for insightful scientific discussions and Prof. Miguel Ángel Silvestre (University of Valencia) for his valuable feedback on the bovine IVF workflow. We also thank the staff at Vorwerk Podemus for assistance with bovine ovary and oviduct sample collection, and M.Sc. Minas Stavrinou for his support to R.N. in the microrobot swarm formation experiments.

## Author Contributions

C.R. and F.S contributed equally to this work. M.M.S. conceived and supervised the study. C.R., F.S and M.M.S designed the experiments. J.S. contributed to the IVF and BOEC sperm binding experiment design. C.R.C, F.S., R.N., F.H. carried out experiments. C.R. and F.S. performed the experimental work and corresponding data collection and analysis. R.N. contributed with the optical feedback control of the magnetotactic sperms. C.R., F.S. and M.M.S. wrote the manuscript with contributions from all authors. All authors revised the manuscript critically before submission.

