## Supporting Information for "In Vitro Fertilization using Magnetotactic Sperm Cells"

**Table S1. Number, average and standard deviation of free sperm and MSC counted for basic purification method with batched 1 of microparticles**

| Group | Number of free sperm | Number of MSC | % free sperm | % MSC |
|---|---|---|---|---|
| **Replicate 1** | 2 | 1315 | 0.15 | 99.84 |
| **Replicate 2** | 5 | 1581 | 0.31 | 99.68 |
| **Replicate 3** | 20 | 1317 | 1.49 | 98.50 |
| **Average** | 9 | 1404.33 | 0.65 | 99.35 |
| **SD** | 9.64 | 153.00 | 0.73 | 0.73 |

**Table S2. Number, average and standard deviation of free sperm and MSC counted for basic purification method with Batched 2 of microparticles**

| Group | Number of free sperm | Number of MSC | % free sperm | % MSC |
|---|---|---|---|---|
| **Replicate 1** | 0 | 341 | 0 | 100 |
| **Replicate 2** | 9 | 995 | 0.90 | 99.10 |
| **Replicate 3** | 11 | 792 | 1.37 | 98.63 |
| **Average** | 6.66 | 709.33 | 0.76 | 99.24 |
| **SD** | 5.89 | 334.75 | 0.70 | 0.70 |

**Table S3. Number, average and standard deviation of free sperm and MSC counted for magnetic separation method with batch 2 of microparticles**

| Group | Number of free sperm | Number of MSC | % free sperm | % MSC |
|---|---|---|---|---|
| **Replicate 1** | 55 | 19084 | 0.29 | 99.71 |
| **Replicate 2** | 45 | 8611 | 0.52 | 99.48 |
| **Replicate 3** | 13 | 12106 | 0.10 | 99.90 |
| **Average** | 37.67 | 13267 | 0.30 | 99.70 |
| **SD** | 21.94 | 5332.16 | 0.21 | 0.21 |

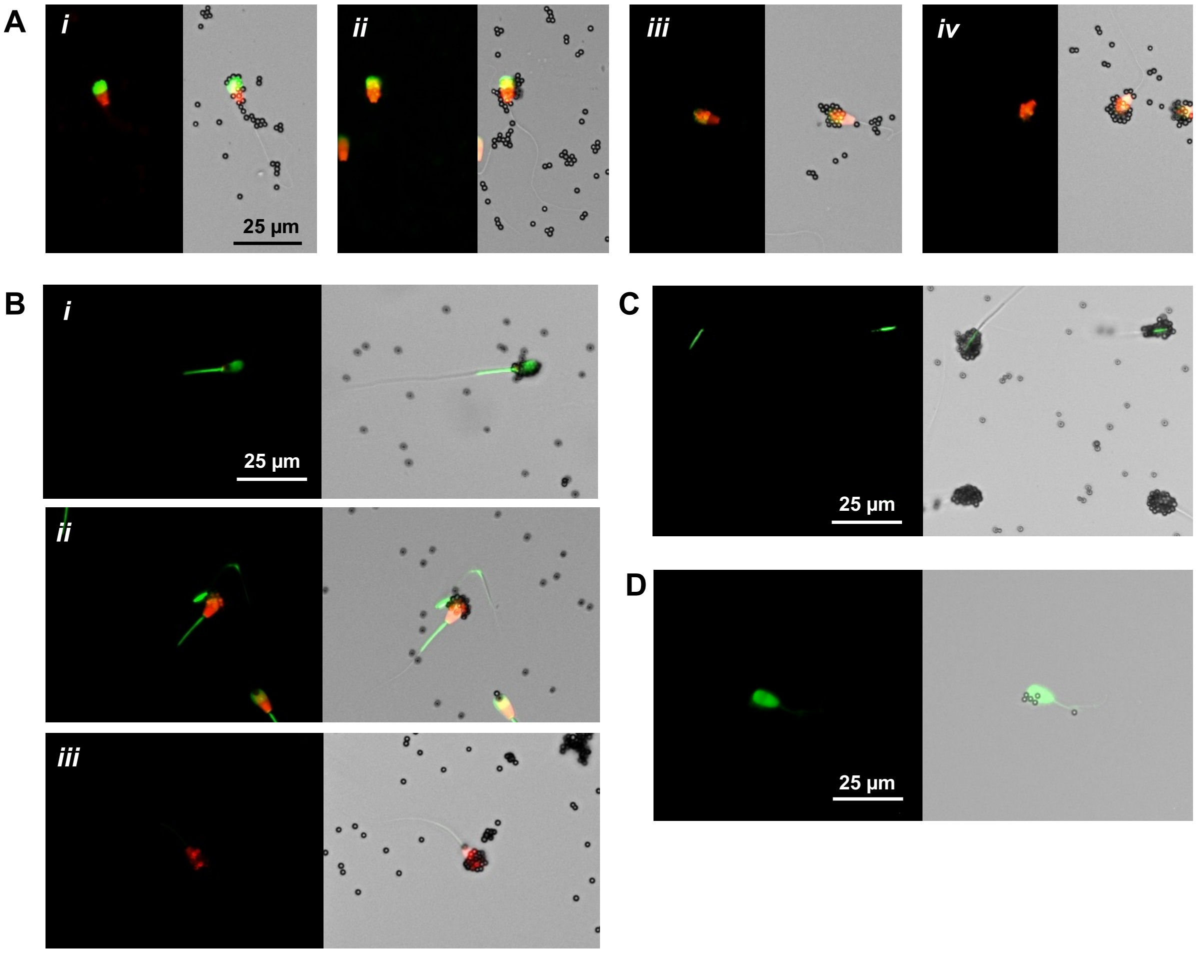


**Figure S1. Representative images of MSC staining for**: A) Acrosome integrity: intact (i), low staining (ii), clearly damaged (iii) and totally lost (iv), B) Mitochondria Membrane Integrity: intact membrane (i), defective membrane (ii) and damaged membrane (iii), C) Oxidative Stress: sperm presenting oxidative stress (top) and non-stress sperm (bottom), and D) DNA fragmentation: sperm with fragmented DNA. Left images of fluorescent filters and right images have fluorescent filters and brightfield.

**Table S4. Acrosome integrity test for free sperm and MSC samples at 0h and 2h incubation**

| | **Free Sperm** | | | | **MSC** | | | |
|---|---|---|---|---|---|---|---|---|
| | **0h** | | | | | | | |
| | **Intact, n (%)** | **Low staining, n (%)** | **Clearly damaged, n (%)** | **Totally lost, n (%)** | **Intact MSC, n (%)** | **Low staining, n (%)** | **Clearly damaged, n (%)** | **Totally lost, n (%)** |
| **Replicate 1** | 781 (59.17) | 312 (23.64) | 177 (13.41) | 50 (3.79) | 185 (79.06) | 21 (8.97) | 11 (4.70) | 17 (7.26) |
| **Replicate 2** | 685 (78.02) | 79 (9.0) | 84 (9.57) | 30 (3.42) | 482 (79.54) | 45 (7.43) | 54 (8.91) | 25 (4.13) |
| **Replicate 3** | 346 (73.77) | 72 (15.35) | 19 (4.05) | 32 (6.82) | 960 (77.61) | 155 (12.53) | 92 (7.44) | 30 (2.43) |

| | | | | | | | | |
|---|---|---|---|---|---|---|---|---|
| **Average** | 604 (70.32) | 154.33 (16.0) | 93.33 (9.01) | 37.33 (4.68) | 542.33 (78.73) | 73.67 (9.64) | 52.33 (7.02) | 24 (4.61) |
| **SD** | 228.53 (9.89) | 136.59 (7.34) | 79.41 (4.70) | 11.01 (1.87) | 391.00 (1.01) | 71.50 (2.62) | 40.53 (2.14) | 6.56 (2.46) |
| **2h** | | | | | | | | |
| **Replicate 1** | 416 (58.67) | 90 (12.69) | 147 (20.73) | 56 (7.90) | 232 (62.87) | 33 (8.94) | 48 (13.01) | 56 (15.17) |
| **Replicate 2** | 263 (51.37) | 49 (9.57) | 70 (13.67) | 130 (25.39) | 347 (52.10) | 58 (8.71) | 85 (12.76) | 176 (26.43) |
| **Replicate 3** | 714 (51.93) | 399 (29.01) | 124 (9.02) | 138 (10.04) | 104 (53.61) | 11 (5.67) | 29 (14.95) | 50 (25.77) |
| **Average** | 464.33 (53.99) | 179.33 (17.09) | 113.67 (14.47) | 108 (14.44) | 227.67 (56.20) | 34 (7.77) | 54 (13.57) | 94 (22.46) |
| **SD (±)** | 229.35 (4.07) | 191.34 (10.44) | 39.53 (5.90) | 45.20 (9.54) | 121.56 (5.83) | 23.52 (1.83) | 28.48 (1.20) | 71.08 (6.32) |

**Table S5. Mitochondria membrane integrity test for free sperm and MSC samples at 0h and 2h incubation**

| | **Free Sperm** | | | **MSC** | | |
|---|---|---|---|---|---|---|
| **0h** | | | | | | |
| | **Intact membrane, n (%)** | **Defective membrane, n (%)** | **Damaged membrane, n (%)** | **Intact membrane, n (%)** | **Defective membrane, n (%)** | **Damaged membrane, n (%)** |
| **Replicate 1** | 247 (44.43) | 149 (26.80) | 160 (28.78) | 116 (35.15) | 206 (62.42) | 8 (2.42) |
| **Replicate 2** | 331 (40.37) | 290 (35.37) | 199 (24.27) | 98 (41.70) | 125 (53.19) | 12 (5.11) |
| **Replicate 3** | 460 (39.66) | 550 (47.41) | 150 (12.93) | 51 (43.97) | 62 (53.45) | 3 (2.59) |
| **Average** | 346 (41.48) | 329.67 (36.53) | 169.67 (21.99) | 88.33 (40.27) | 131 (56.35) | 7.67 (3.37) |
| **SD (±)** | 87.60 (2.57) | 166.09 (10.36) | 21.14 (8.16) | 27.40 (4.58) | 58.94 (5.26) | 3.68 (1.50) |
| **2h** | | | | | | |
| **Replicate 1** | 220 (25.06) | 273 (31.09) | 385 (43.85) | 85 (44.50) | 98 (51.31) | 8 (4.19) |
| **Replicate 2** | 122 (20.03) | 102 (16.75) | 385 (63.22) | 128 (43.54) | 154 (52.38) | 12 (4.08) |
| **Replicate 3** | 142 (16.06) | 54 (6.11) | 688 (77.83) | 95 (35.19) | 143 (52.96) | 32 (11.85) |
| **Average** | 161.33 (20.38) | 143 (17.98) | 486 (61.63) | 102.67 (41.08) | 131.67 (52.23) | 17.33 (6.71) |
| **SD (±)** | 42.28 (4.51) | 93.99 (12.54) | 142.84 (17.04) | 18.37 (5.12) | 24.22 (0.84) | 10.50 (4.46) |

**Table S6. Oxidative stress test for free sperm and MSC samples at 0h and 2h incubation**

| | Free sperm | | MSC | |
|---|---|---|---|---|
| **0h** | | | | |
| | **Stress, n (%)** | **Not stressed, n (%)** | **Stress, n (%)** | **Not stressed, n (%)** |
| **Replicate 1** | 14 (3.94) | 341 (96.06) | 3 (0.63) | 477 (99.38) |
| **Replicate 2** | 12 (4.84) | 236 (95.16) | 7 (1.19) | 581 (98.81) |
| **Replicate 3** | 13 (5.44) | 226 (94.56) | 1 (0.31) | 325 (99.69) |
| **Average** | 13 (4.74) | 267.67 (95.26) | 3.67 (0.71) | 461 (99.29) |
| **SD (±)** | 1 (0.75) | 63.71 (0.75) | 3.06 (0.45) | 128.75 (0.45) |
| **2h** | | | | |
| **Replicate 1** | 34 (11.07) | 273 (88.93) | 9 (1.27) | 702 (98.73) |
| **Replicate 2** | 5 (1.85) | 266 (98.15) | 2 (0.39) | 505 (99.61) |
| **Replicate 3** | 35 (10.39) | 302 (89.61) | 2 (0.40) | 501 (99.60) |
| **Average** | 24.67 (7.77) | 280.33 (92.23) | 4.33 (0.69) | 569.33 (99.31) |
| **SD (±)** | 17.04 (5.14) | 19.09 (5.14) | 4.04 (0.50) | 114.91 (0.50) |

**Table S7. Sperm DNA fragmentation test for free sperm and MSC samples at 0h and 2h incubation**

| | Free sperm | | MSC | |
|---|---|---|---|---|
| **0h** | | | | |
| | **Fragmented, n (%)** | **Non fragmented, n (%)** | **Fragmented, n (%)** | **Non Fragmented, n (%)** |
| **Replicate 1** | 83 (2.39) | 3389 (97.61) | 7 (2.48) | 275 (97.52) |
| **Replicate 2** | 7 (0.21) | 3288 (99.79) | 2 (0.95) | 207 (99.04) |
| **Replicate 3** | 29 (1.71) | 1662 (98.29) | 0 (0) | 163 (100) |
| **Average** | 39.67 (1.44) | 2779.67 (98.56) | 3 (1.15) | 215 (98.85) |
| **SD (±)** | 39.1 (1.11) | 969.24 (1.11) | 3.61 (1.25) | 56.43 (1.25) |
| **2h** | | | | |
| **Replicate 1** | 8 (0.37) | 2128 (99.63) | 1 (1.35) | 73 (98.65) |
| **Replicate 2** | 2 (0.08) | 2497 (99.92) | 9 (3.67) | 236 (96.33) |
| **Replicate 3** | 9 (0.32) | 2836 (99.68) | 1 (0.38) | 258 (99.61) |
| **Average** | 6.33 (0.26) | 2487 (99.74) | 3.67 (1.80) | 189 (98.20) |
| **SD (±)** | 3.79 (0.16) | 354.11 (0.16) | 4.62 (1.69) | 101.06 (1.69) |

**Table S8. Number of free sperm and MSC attached to BOECs ciliated cells and the location attachment in the sperm head**

| Ciliated cells attachment | Free sperm, n (%) | MSC, n (%) |
|---|---|---|
| **Acrosome** | 12 (52.17) | 14 (35) |
| **Head** | 11 (47.83) | 26 (65) |

**Table S9. IVF results across different experimental groups in two replicates**

| | Total COCs, n | Immature oocytes, n | Maturation rate, n (%) | Cleavage rate, n (%) | Blastocyst rate (Blastocyst/Cleavage embryos), n (%) | Blastocyst rate (Blastocyst/Mature oocytes), n (%) |
|---|---|---|---|---|---|---|
| **IVF 1** | | | | | | |
| **P** | 25 | 4 | 21 (84) | 0 (0) | 0 (0) | 0 (0) |
| **P + P** | 33 | 6 | 27 (82) | 0 (0) | 0 (0) | 0 (0) |
| **C.IVF** | 36 | 2 | 34 (94) | 25 (74) | 14 (56) | 14 (41) |
| **MC** | 39 | 5 | 34 (87) | 5 (15) | 0 (0) | 0 (0) |
| **MSC** | 33 | 6 | 27 (82) | 6 (22) | 2 (33) | 2 (7) |
| **LC** | 35 | 2 | 33 (94) | 1 (3) | 0 (0) | 0 (0) |
| **IVF 2** | | | | | | |
| **P** | 60 | 7 | 53 (88) | 8 (15) | 0 (0) | 0 (0) |
| **P + P** | 53 | 7 | 46 (87) | 0 (0) | 0 (0) | 0 (0) |
| **C.IVF** | 53 | 9 | 44 (83) | 35 (80) | 18 (51) | 18 (41) |
| **MC** | 52 | 1 | 51 (98) | 7 (14) | 4 (57) | 4 (8) |
| **MSC** | 51 | 4 | 47 (92) | 7 (15) | 3 (43) | 3 (6) |
| **LC** | 49 | 4 | 45 (92) | 1 (2) | 0 (0) | 0 (0) |

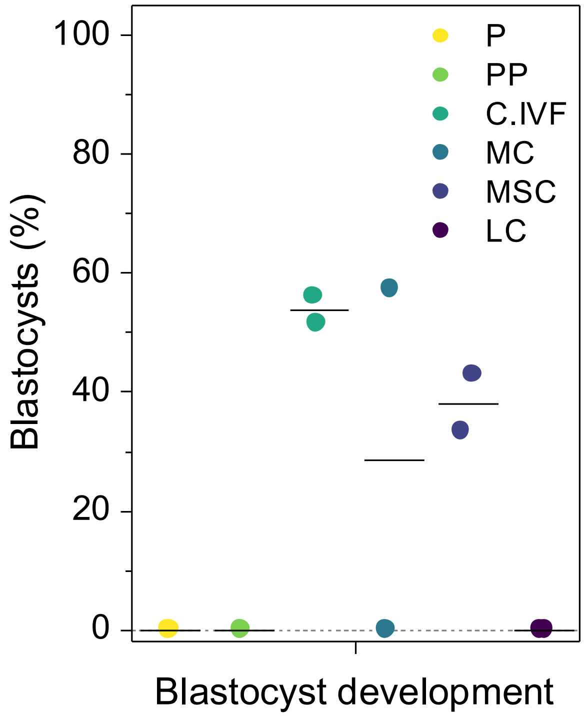


**Figure S2. Blastocyst formation from cleavage stage embryos.**

**Table S10. Blastocyst and hatched blastocyst formation across different experimental groups on D9 (two replicates)**

| | P | P + P | C.IVF | MC | MSC | LC |
|---|---|---|---|---|---|---|
| **IVF 1** | | | | | | |
| **Blastocyst, n (%)** | 0 (0) | 0 (0) | 4 (28.57) | 0 (0) | 0 (0) | 0 (0) |
| **Hatched blastocyst, n (%)** | 0 (0) | 0 (0) | 10 (71.43) | 0 (0) | 2 (100) | 0 (0) |
| **IVF 2** | | | | | | |
| **Blastocyst, n (%)** | 0 (0) | 0 (0) | 5 (27.78) | 1 (25) | 1 (16.67) | 0 (0) |
| **Hatched blastocyst, n (%)** | 0 (0) | 0 (0) | 13 (71.83) | 3 (75) | 2 (83.33) | 0 (0) |
